# Detection of *Deltacoronavirus* in Environmental Fecal Samples from Seabirds in the Saint Peter and Saint Paul Archipelago, central equatorial Atlantic Ocean

**DOI:** 10.1101/2024.11.19.624433

**Authors:** Fernanda Gomes, Alexandre Freitas da Silva, Tatiana Prado, Marina Galvão Bueno, Luciana Appolinario, Patrícia Soares Flores, Paola Cristina Resende, Marilda Siqueira, Leonardo Corrêa, Martha Brandão, Jose Reck, Gabriel da Luz Wallau, Maria Ogrzewalska

## Abstract

This study investigates the presence of avian coronaviruses (CoVs), influenza A viruses (IAVs), and Avian rotaviruses (RVs) group A in seabird populations inhabiting the Saint Peter and Saint Paul Archipelago (SPSPA), an isolated and remote oceanic island situated in the equatorial region of the Atlantic Ocean. In July 2022, 95 environmental fecal samples were collected from seabird colonies and screened for the viruses by quantitative one-step real-time RT-PCR (IAVs and RVs), by the conventional pancoronavirus RT-PCR protocols and metatranscriptomics of a positive sample. Four environmental feces samples tested positive for CoVs. Avian IAVs and RVs were not detected. Phylogenetic analysis revealed CoVs closely related to avian deltacoronaviruses previously identified in waterbirds from Asia and Australia. We could not recover the CoV by metatranscriptomics but we recovered a single viral contig of an avian enterovirus. The findings contribute valuable insights into virus dynamics among seabird populations, laying the groundwork for future investigations in this field.

## Introduction

Seabirds comprise a vast diversity of species and include members of at least six avian orders such as Sphenisciformes, Procellariiformes, Pelecaniformes, Suliformes, Phaethontiformes and Charadriiformes, that all share the characteristic of feeding at sea (Furness, 1987). A vast majority of seabirds exhibit colonial breeding behavior, gathering in large numbers for extended periods each year to reproduce and although seabirds exhibit remarkable colony fidelity for breeding, they are also known to undertake extensive journeys during their non-breeding periods to forage and explore potential future breeding sites (Furness, 1987). This behavior highlights the potential role of seabirds as important reservoirs in the dispersal of infectious agents across diverse ecosystems (1-4).

The Saint Peter and Saint Paul Archipelago (SPSPA) is a small group of rocky islets and reefs of volcanic origin, located in the middle of the Atlantic Ocean. Only three seabird species reproduce in the SPSPA: Black noddy *Anous minutus* (Boie, F., 1844), Brown noddy *Anous stolidus* (Linnaeus, 1758) (Charadriiformes, Laridae) and Brown booby *Sula leucogaster* (Boddaert, 1783) (Suliformes, Sulidae). Noddies breed here between March and September(5), while Brown boobies breed throughout the year, showing the absence of seasonality in nesting (5, 6). Information on the migratory behavior of noddies and bobbies at the SPSPA remains limited. However, a few studies suggest that these populations are not migratory within the region. This conclusion is based on observations that their population sizes do not exhibit significant seasonal variations throughout the year, nor is there evidence of substantial migrations involving large portions of the population (6). Nonetheless, there is a risk of interspecies viral transmission, as the SPSPA is sporadically visited by various seabird species such as masked booby *Sula dactylatra* (Lesson, 1831), red-footed booby *Sula sula* (Linnaeus, 1766), magnificent frigatebird *Fregata magnificens* (Mathews, 1914) (Suliformes, Fregatidae), sooty tern *Onychoprion fuscatus* (Linnaeus, 1766) (Charadriiformes, Laridae). Other migratory water birds are known to appear, usually lost during their migration movements, such as herons; cattle egret *Bubulcus ibis* (Linnaeus, 1758) (Pelecaniformes, Ardeidae), little egret *Egretta gularis* (Bosc, 1792) and white egret *Egretta garzetta* (Linnaeus, 1766), lesser moorhen *Paragallinula angulata* (Sundevall, 1851) (Gruiformes, Rallidae) and even birds of prey such as falcon *Falco tinnunculus* (Falconiformes, Falconidae) and shorebirds turnstone *Arenaria interpres* (Linnaeus, 1758) (Charadriiformes Scolopacidae) and yellowlegs *Tringa flavipes* (Gmelin, JF, 1789) (7). The archipelago is inhabited, with no domestic animals or wild mammals present. However, the largest island, Belmonte Island, hosts a Brazilian Navy research station. This station maintains a full-time presence of researchers working on various scientific projects (8).

Research on seabird parasites and pathogens has suggested a diverse range of infecting organisms associated with these birds including ectoparasites, protozoa, bacteria, and viruses (2, 9-14). Among viruses associated with seabirds, coronaviruses (CoVs), influenza A viruses (IVAs) and avian rotaviruses (RVs) warrant special attention due to their potential to cause morbidity and in some cases mass mortality among both wild and domestic birds. These viruses can have significant impacts on avian populations, leading to severe outbreaks and affecting the health of individual birds as well as the overall dynamics of bird communities (15-22). No previous study has been conducted in this area to investigate the presence of viruses. To address this research gap, we conducted a study focusing on collecting environmental samples from seabirds to test for the presence of CoVs, IVAs and group A RVs. By examining these samples, we aimed to gain valuable insights into the potential role of seabirds as hosts of these viruses in this unique and isolated ecosystem.

## Materials And Methods

### Study area and sample collection

The SPSPA is part of the territorial waters of Brazil and is situated approximately 1 010 km off the northeastern coast of Brazil (00°55’10"N, 29°20’33"W). The rocky formation of the Archipelago covers an area of approximately 17 000 m^2^, with a maximum altitude of 18 m. The SPSPA consists of six main islets: Cabral, Nordeste, Challenger, Belmonte, South and Coutinho (Figure 1).

**Figure 1.**
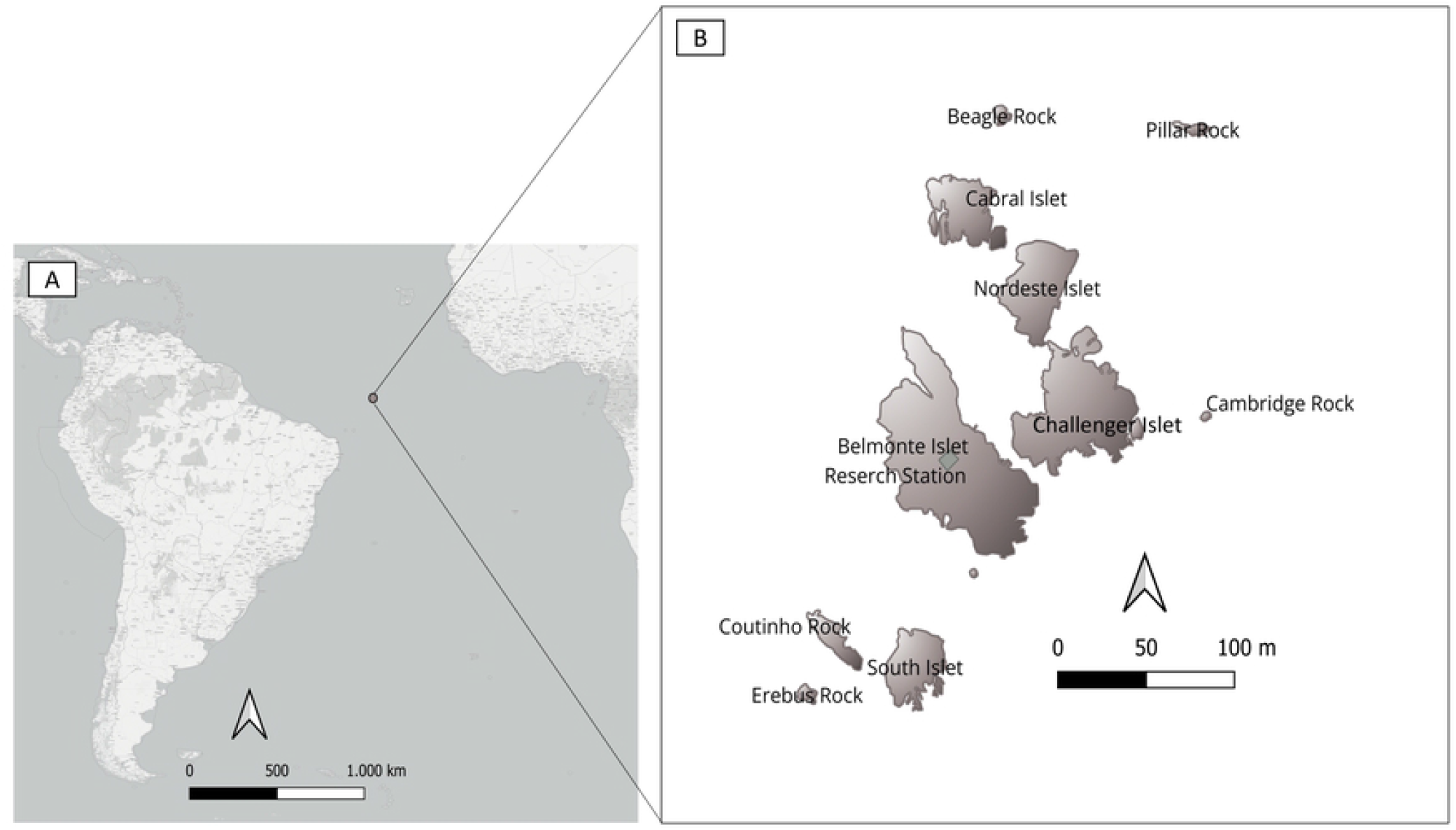
Localization of Saint Peter and Saint Paul Archipelago, central equatorial Atlantic Ocean (A). Saint Peter and Saint Paul Archipelago with main islets, rocks, and research station (B). Map created using the Free and Open Source QGIS.

**Figure 2.**
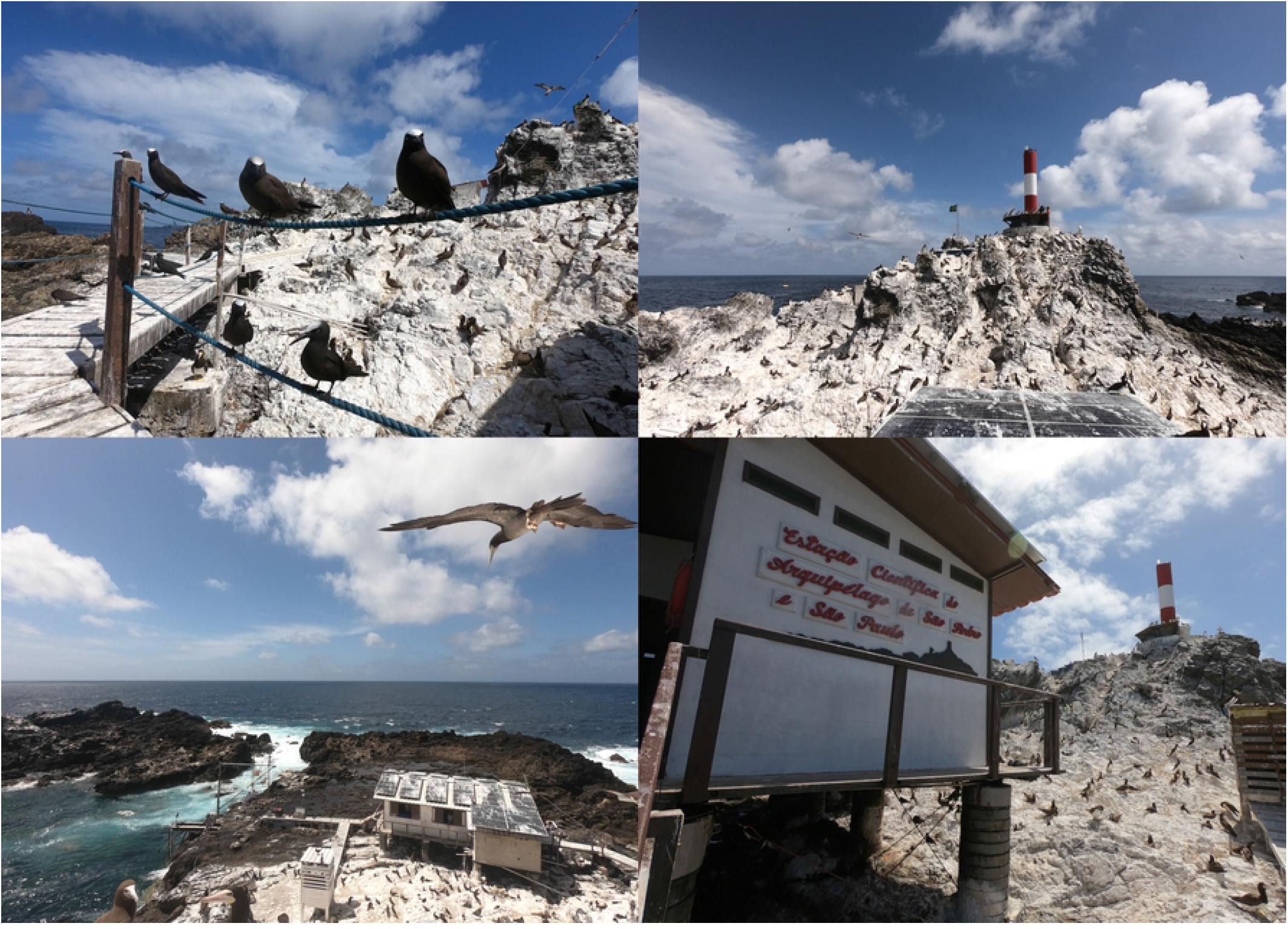
Images from the Baltimore Island show high densities of seabirds and the research station covered in guano. July 2022, Saint Peter and Saint Paul Archipelago, central equatorial Atlantic Ocean. Photos by M. Brandao.

Convenience fresh environmental fecal samples were collected in July 2022 on the Belmonte islet, during three different consecutive days. Fresh dropping samples were collected using sterile Dacron swabs, which were immediately placed into sterile tubes containing 1 mL of RNA*later*™ Stabilization Solution (Invitrogen™). The samples were kept at room temperature for seven days before being frozen at −20°C for further analysis.

### Viral RNA extraction

Clarified fecal suspensions (20%, wt/vol) were prepared with phosphate-buffered saline (PBS) as previously described (23). Viral RNA was extracted using a QIAamp viral RNA mini kit (Qiagen, CA, USA) according to the manufacturer’s instructions. For each RNA extraction procedure, RNase/DNase-free water was used as a negative control.

### Influenza A virus (IAVs) and group A avian rotavirus (AvRVA) screening by quantitative one-step real-time RT-PCR

The IAVs and AvRVA were screened using TaqMan based quantitative one-step real-time RT-PCR designs. All reactions were performed using the SuperScript III Platinum one-step quantitative RT-PCR (qRT-PCR) kit (Thermo Fisher Scientific) according to the protocol established by the Collaborative Influenza Center, Centers for Disease Control and Prevention, Atlanta, GA for detection of all influenza A (24) and (25) for detection of all rotavirus A. The samples with Cycle threshold (Ct) < 40 were considered positive.

### Coronavirus (CoVs) screening by conventional pancoronavirus RT-PCR

All the samples were also subjected to pancoronavirus RT-PCR targeting the RNA-dependent RNA polymerase (*RdRp*) gene as described previously (26). Briefly, cDNA was obtained and amplified in a first-round PCR (RdRpS1 5’-GGKTGGGAYTAYCCKAARTG-3’, RdRpR1 5’-TGYTGTSWRCARAAYTCRTG-3’) using One-Step RT-PCR Enzyme MixKit (Qiagen) with the total expected size of 602 base pairs (bp). Following, the nested PCR were conducted using Phusion RT-PCR Enzyme Mix kit (Sigma-Aldrich), primers Bat1F 5’-GGTTGGGACTATCCTAAGTGTGA-3’ and Bat1R 5’-CCATCATCAGATAGAATCATCAT-3’ and 1 uL of the amplified product as a template were used. *RdRp* amplicons (∼440 bp) were visualized on 1.5% agarose gels with SYBR™ Safe DNA Gel Stain (Thermo Fisher Scientific).

### Sanger sequencing

*RdRp* amplicons (440 bp) were purified using the QIAquick Gel Extraction Kit (Qiagen) following the manufacturer’s recommendation. The Sanger sequencing reaction was prepared using BigDye Terminator v3.1 Cycle Sequencing Kit (Life Technologies) with primers Bat1F and Bat1R. The sequencing of both strands was performed using the ABI 3730 DNA Analyzer (Applied Biosystems).

### Gene assembly and phylogenetic analysis

The reads generated by Sanger sequencing were evaluated and assembled using Sequencher 5.1 (GeneCodes). The obtained consensus was checked by the chromatogram analysis and the final consensus was uploaded at the National Center for Biotechnology Information (NCBI) GenBank database. To enhance our understanding of the phylogenetic relationships among the sequences generated in this study we performed homologous sequence retrieval from the non-redundant NCBI database based on blastn of all partial RdRp obtained in this study. The dataset for analysis was compiled to include: (i) sequences determined in this work (n = 3), (ii) the most similar sequences from GenBank to those from this work, obtained from a BLAST analysis (hits with the highest score, identity > 70%, and coverage > 60%, excluding duplicates, n = 54), (iii) reference sequences for all three subgenera of the genus *Deltacoronavirus* as defined by the ICTV (n = 7), and (iv) reference sequences from members of the genus *Gammacoronavirus* (n = 6) for rooting the tree (ICTV, 2022). Nucleotide alignments were performed with MAFFT v7.0 online (https://mafft.cbrc.jp/alignment/server/index.html). (https://ictv.global/report/chapter/coronaviridae/coronaviridae). The best nucleotide substitution model was defined by MEGA X and used for the reconstruction of the maximum likelihood (ML) phylogenetic tree of the partial *RdRp* gene (27) with bootstrap values obtained with 500 replicates. The ML tree was annotated in FigTree v.1.4.4 (http://tree.bio.ed.ac.uk/software/figtree/).

### Metatranscriptomic sequencing

One CoV positive sample (A22801) that reached enough RNA quantity and quality was processed to metatranscriptomic sequencing aiming to characterize their viral content. The previously extracted RNA was treated with Ambion® TURBO DNA-free™ Kit (Invitrogen) following the manufacturer’s instructions to remove residual genomic DNA. Host rRNA was depleted using the Illumina Ribo-Zero Plus rRNA Depletion Kit (Illumina, San Diego, CA, USA). After rRNA depletion, the remaining RNA was converted into cDNA and prepared for library construction using the Illumina® DNA Prep Kit. The libraries were sequenced on the Illumina® NextSeq platform (Illumina, San Diego, CA, USA) using a NextSeq 1000/2000 P2 cartridge employing a paired-end approach. The raw submitted to NCBI sequence Read Archive (SRA) database and are available under project number: PRJNA1183304.

After sequencing the raw reads were first analyzed on fastp v0.23.2 and the low-quality reads were trimmed with a cutoff of 20 for Phred score and a minimum read length of 36 bp. Trimmed reads were assembled using a *de novo* approach on metaSPAdes v3.15.5 in default mode (28). The assembled contigs were submitted to a pairwise-alignment analysis employing Diamond blastx (29) against a custom dataset constructed based on all viral proteins assigned to taxonomy tag (txid10239) from NCBI plus RdRp sequence databases such as NeoRdRp (30), PalmDB (31) and RdRp-Scan (32) as performed previously by (33). In brief, the sequences showing the best hits on the viral database were submitted to a second Diamond blastx analysis against the non-redundant (NR) database from NCBI to remove false negative hits. We also performed a BLASTn (34) analysis to identify the best nucleotide hits based on the NT database from NCBI. The completeness of viral sequences were assessed on ViralComplete (35) tool that classified them into full-length or partial according to their best hit with NCBI RefSeq genomes.

All applicable national and institutional guidelines for sampling, care and experimental use of organisms for the study have been followed and all necessary approvals have been obtained such as Permits of Instituto Chico Mendes de Conservação da Biodiversidade (SisBio 84084-1) and the registration of the project in the Sistema Nacional de Gestão do Patrimônio Genético e do Conhecimento Tradicional Associado (A21FE59).

All permits are available on request.

## Results

During three consecutive days of collecting fresh droppings, only three bird’s species were observed on the islands: *A. minutus, A. stolidus* and *S. leucogaster*. A total of 95 environmental fecal samples were collected and tested for selected viruses. Four samples tested positive for CoVs targeting *RdRp* genome region. After the *RdRp* nucleotide Sanger sequencing of the positive samples it was possible to obtain fragments of genomes from three of them. The three *RdRp* fragment sequences are available at GenBank database as AvCoV/env/Brazil/PE/FIOCRUZ-A220778/2022 (OR344774), AvCoV/env/Brazil/PE/FIOCRUZ-A220801/2022 (OR344775) and AvCoV/env/Brazil/PE/FIOCRUZ-A220810/2022 (OR344776). Obtained viruses were 99.5-99.7% identical between themselves and the closest identity of 97% (218/225 bp) was observed with avian *Deltacoronavirus* found in swan goose *Anser cygnoides* (Linnaeus, 1758) (Anseriformes, Anatidae) from southern Brazil (KU321643, KU321644) in 2013. The phylogenetic reconstruction of *RdRp* gene fragment revealed the three CoVs detected in the present study cluster in the unnamed clade of deltacoronaviruses, with a sister clade of viruses detected previously in herons (Pelecaniformes, Ardeidae) from Vietnam and Australia in 2020 and 2016 respectively (Figure 3). The sequences from Brazil were not included in phylogenetic analyses because the deposited in GenBank RdRp gene fragments were only 225 bp long, with coverage of 54% with our sequences.

**Figure 3.**
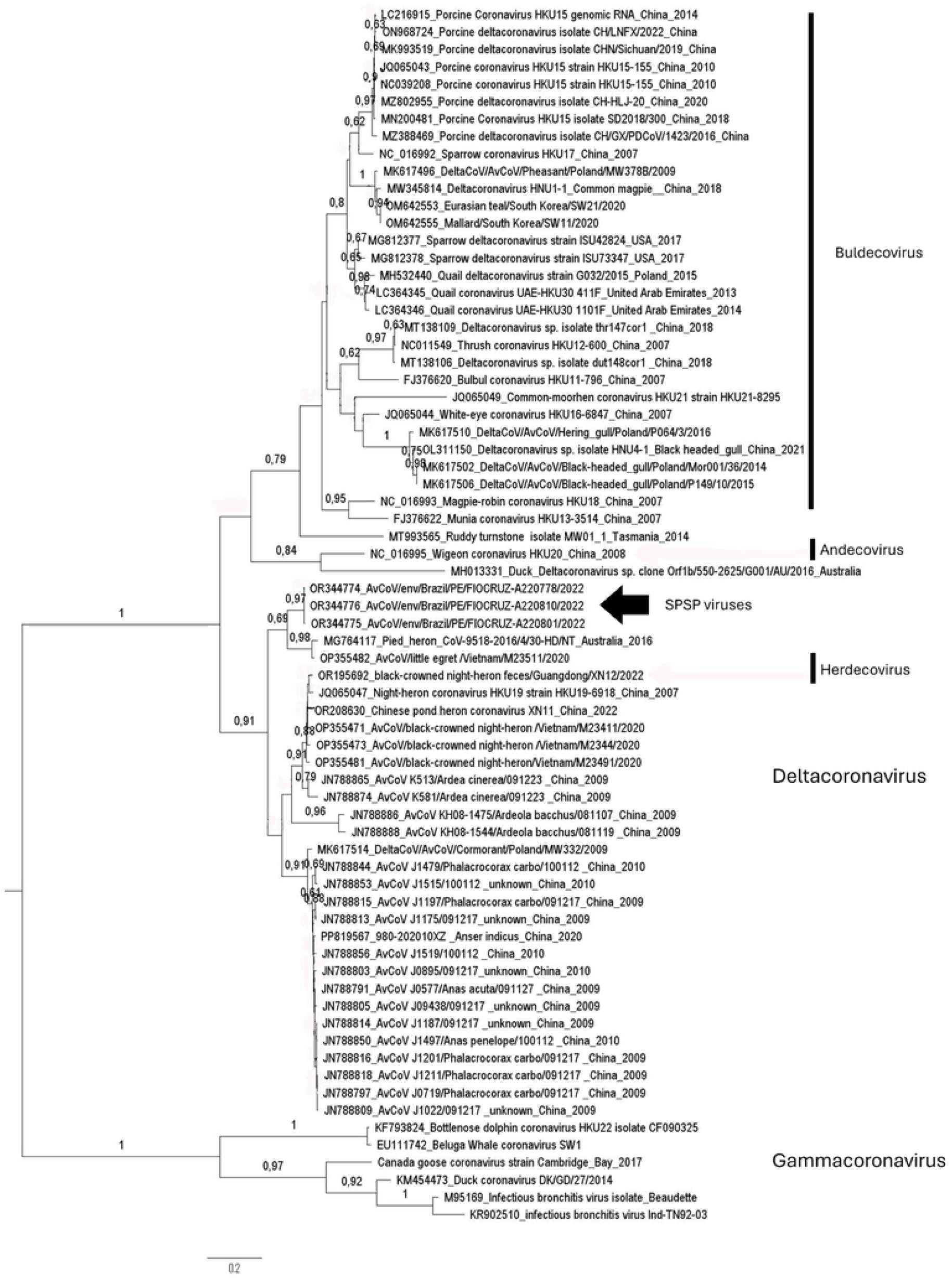
A Maximum Likelihood phylogenetic tree, focusing on deltacoronaviruses, was reconstructed using the best-fit nucleotide substitution model, General Time Reversible (GTR+G). This tree highlights the positions of viruses identified in the present study (arrow). The final dataset comprised 70 partial genomes, specifically 360 base pairs of the RdRp gene.

In addition to our previous analysis, we further investigated one sample positive for CoV (A22801) using a metatranscriptomic approach. The sequencing yielded 57.28 million raw reads and 50.72 million quality-filtered reads were assembled. Diamond analysis revealed the presence of three viral Enterovirus contigs with size ranging from 1,494 to 3,352 bp but no Deltacoronavirus contigs were detected. Enterovirus contigs showed high identity with human Enterovirus 99, with amino acid identities ranging from 52% to 98.5 (ABM54518.1 and QRG33086.1) and nucleotide analysis identities ranging from 82.7% to 84.06 (EF015008.1 and EF015009.1).

Avian IAVs and RVs were not detected in all tested samples.

## Discussion

Most deltacoronaviruses have been found in birds, the only exception being porcine coronavirus HKU15, which is found in pigs and occasionally in humans. Deltacoronaviruses have been identified across 15 bird orders, involving over 100 species of wild birds. They are most frequently observed in waterfowl and shorebirds, particularly within the Charadriiformes and Anseriformes orders (26, 36, 37). Three subgenera can be distinguished within the *Deltacoronavirus* genus, namely *Andecovirus* (one species), *Buldecovirus* (five species) and *Herdecovirus* (one species). However, we were not able to classify the viruses found in the present study into any of these three subgenera. We acknowledge the limitations of our work, as we were unable to recover the full genome of the obtained viruses and conduct more detailed analyses. Our viruses, as well as many others available in GenBank, present only small genome fragments and only viruses for which a complete genome sequence are available might be considered for taxonomy to define possible new subgenera (38).

Previous research has shown that wild aquatic birds of the order Anseriformes and Charadriiformes are the natural reservoir of AIVs (39). However, our analysis did not reveal the presence of influenza virus in any of the collected environmental samples. This outcome contrasts with previous studies reporting positive detections of avian influenza viruses in similar habitats (19). The absence of influenza virus detection in the collected environmental samples could be attributed to several factors and does not necessarily imply an absence of the virus itself. The short duration of the sampling period, which was limited to only three days, might have influenced the absence of influenza virus detection. Our brief three-day sampling window might not have encompassed the entirety of these shedding patterns, potentially missing windows of active viral shedding. Additionally, a limited number of samples was collected and tested.

Just like the absence of influenza virus infection, the lack of detection of Avian RVA in the feces of the birds can be explained by short sampling period and the possibility of prior acquisition of immunity, considering the long-life span of the birds. However, it could be assumed that there is no circulation of AvRVA in the bird populations that inhabit the island, in the period sampled, since fresh feces were collected and the high capacity of these viruses to survive in the environment. *Rotavirus* are non-enveloped viruses, which makes them stable and infective for a long period in the environment without suffering degradation and inactivation. As well as other enteric viruses, all RVAs are resistant to chemical and physical agents in the environment (40). RVAs are not well studied among wild-living birds including seabirds, however, some recent studies suggest that the genetic diversity of avian RVAs is greater than previously recognized, infections occur in a wide spectrum of bird species and that migratory birds may contribute to the global spread of these viruses (41-43).

The ASPSP archipelago is regarded as one of the most remote and isolated locations in the Atlantic Ocean. Although not inhabited by humans, it receives periodic visits from Brazilian Navy personnel who maintain a small research station on one of the islets. This station serves as a hub for scientific research, with researchers residing on the ASPSP for extended periods, often spanning several days or even weeks (8). In this time, researchers share the same habitat as marine birds, which puts their health at risk due to the ubiquitous presence of bird guano. It is worth emphasizing the potential hazards associated with this situation, including the transmission of pathogens to humans. This concern becomes particularly relevant considering the ongoing spread of the highly pathogenic subtype of bird flu H5N1 across South America for the first time that affects not only birds but mammals including humans as well (18, 44-46). Migrant species stopping at Archipelago could potentially bring the H5N1 virus from distant regions introducing it to the area. This situation poses a risk not only to the resident bird population but also to human health.

## Conclusion

This study shows that marine birds from the ASPSP are important in the circulation of new variants of avian coronaviruses, with unknown pathogenicity. Understanding the prevalence of pathogens in wild birds is important for assessing their role in the overall epidemiology of these pathogens, including potential spillover events to domestic animals and humans. Continued and enhanced surveillance efforts are essential to monitor the presence of viruses, such as CoVs, IVAs, and avian RVs group A in avian populations and their potential threat to human and animal health.

## Acknowledgements

Rede de Plataformas Tecnológicas Fiocruz, Instituto Oswaldo Cruz, Fundação de Amparo à Pesquisa do Estado do Rio Grande do Sul (FAPERGS) and Programa Inova FIOCRUZ. Brazilian Navy and Secretaria da Comissão Interministerial para os Recursos do Mar (SECIRM) for logistical support and transportation to the archipelago. Special thanks to Captain Marco Antonio Carvalho de Souza and Sandra Pereira Soares from Fiocruz and finally we thank the anonymous reviewers.

